# Phenological and epidemiological impacts of climate change on peach production

**DOI:** 10.1101/2023.11.13.566906

**Authors:** Chiara Vanalli, Andrea Radici, Renato Casagrandi, Marino Gatto, Daniele Bevacqua

## Abstract

Agricultural food security is threatened by climate change impacts, which can distress crop growth and favor the spread of infectious diseases. Here, we examined the synergism of the two potentially most disruptive causes of future yield failure in peach production: the effects of global climate change on fruit growth and on the spread of fungal diseases. Coupling a phenological and epidemiological model across the French continental territory, we provided projections of yield losses for four peach cultivars (early, mid-early, mid-late, and late) in the XXI century under different climate change scenarios. Global warming is expected to impair fruit phenology with blooming failure events in the south-western part of the country. This will be less extreme under the more moderate emission scenario, even though sporadic blooming failures will still occur. In contrast, future warmer and drier conditions will decrease brown rot-induced yield loss in the historical locations devoted to peach cultivation. To adapt to these changes, the benefits of shifting peach production sites to new suitable areas are evaluated. Thanks to this strategy, the peach national yield could still be fulfilled even under the most extreme climate change scenario. Comprehensive mathematical frameworks, that concomitantly consider the climatic effects on the plant hosts and on their pathogens, are required to provide reliable future predictions of crop yields and to inform control and adaptation strategies to guarantee food security under global warming.

## 1 Introduction

Impacts of climate change include alteration of ecological processes, higher ecosystem vulnerability and shifts in phenology and spatial distribution of species (Pacifici et al., 2015; Hartmann et al., 2022; Walther et al., 2002; Thuiller et al., 2005; Schröter et al., 2005; Parmesan, 2006). From an agricultural point of view, climate change has already been reducing global crop production by 1–5 % per decade over the last years (Porter et al., 2014; Ray et al., 2019). The alarming disparity between reduced food production and population growth is expected to increase in the future (Challinor et al., 2014; Ray et al., 2019), making climate change a threat to global food security. The impacts of climate change on crop yields can be either direct, by impairing plant growth, or indirect, by creating favorable conditions for crop diseases and pests (Chaloner et al., 2021; Newbery et al., 2016; Bebber et al., 2013; Wheeler and Von Braun, 2013). A deeper and comprehensive understanding of these climatic effects on the crops and on their pathogens is therefore urged, and the design and implementation of optimal control and adaptation strategies is required to move food production towards a system more resilient to climate change (Wheeler and Von Braun, 2013; Chakraborty and Newton, 2011).

Temperature is an important driver of perennial plant phenology, determining key events of the annual cycle such as leaf unfolding, dormancy, flowering and fruit ripening (Piao et al., 2019; Chmielewski et al., 2004; Estrella et al., 2007; Hänninen and Tanino, 2011). As a consequence of this dependency, temperature is one of the key factors that shapes the ecological niche of each plant species, characterizing the suitable areas of the globe where the fulfillment of plant thermal requirements is achieved (Vanalli et al., 2021; Menzel and Fabian, 1999; Root et al., 2003; Körner and Basler, 2010; Wolkovich et al., 2017). Recent studies have identified the chilling requirement for dormancy as the most critical phenological process in the future, because it may delay spring phenology or even prevent flowering (Vanalli et al., 2021; Hänninen and Tanino, 2011; Yu et al., 2010; Laube et al., 2014). For fruit orchards, besides phenology, climate and its change have been proved to affect fruit growth and final yield quantity and quality. Warmer temperatures during the growing season may increase fruit growth rate, resulting in an earlier harvest timing (see Tourre et al. (2011) for grapes, Vanalli et al. (2021) for peaches and Warrington et al. (1999) for apples). In addition to temperature, precipitation and water availability are also crucial environmental factors for yield production: extended droughts limit the growth of crops, resulting in a decline of the crop size and quality (Alqudah et al., 2011; Zhang et al., 2018). Future combined effects of population growth and climate change could potentially have detrimental effects on agricultural productivity due to lack of water availability for irrigation (Elliott et al., 2014).

Plant pathogens are extremely diverse and span from intracellular viruses and bacteria to extracellular pathogens, including oomycetes, nematodes and fungi (Velásquez et al., 2018). The latter are responsible for two thirds of the crop losses worldwide (Fisher et al., 2012) and are a paradigmatic case of climate dependency (Gange et al., 2007; Romero et al., 2021). The results of several lab experiments and field observations indicate that the effects of climatic conditions on fungal infections are multiple: i) spore dispersal and release are facilitated by precipitation events (Caubel et al., 2012); ii) spore survival decreases with increasing temperature (Caubel et al., 2012; Xu and Robinson, 2000; Bevacqua et al., 2023); iii) spore penetration into the host tissue positively depends on the fruit surface wetness and on precipitation events (Huber and Gillespie, 1992; Bevacqua et al., 2023); iv) an optimal range of temperatures is needed for the fungal disease to develop (Phillips, 1982; Biggs et al., 1988; Xu et al., 2001a; Chaloner et al., 2021).

Given the described complexity of the climatic dependence of the plant-pathogen systems, a novel framework that embraces the interconnection between these ecological relations is required. With this aim and in light of the new challenges that climate change is posing to agriculture, we propose to implement a well-established framework in plant pathology: the so-called ‘disease triangle’ (Stevens, 1960). According to Stevens Stevens (1960), the environment, the host, and the pathogen must be considered as three components of one interconnected triangular system, with the success of an infection resulting from the interaction between a favorable environment, a susceptible host and a virulent pathogen. Conversely, disease spreading is prevented upon elimination of any one of these fundamental components (Stevens, 1960; Haggag et al., 2016). Evaluating climate change effects concurrently on the plant host, the pathogen and their interactions, it is possible to produce meaningful and realistic future projections of disease spread and crop production.

The evaluation of climate change effects on the plant-pathogen-environment triangle faces numerous challenges. The responses to climate are usually non-linear, making the impacts of environmental changes rarely straightforward (Huey and Kingsolver, 1989; Angilletta Jr et al., 2002). Multiple scenarios of climate change exist, which correspond to different CO_2_ atmospheric concentration pathways (IPCC, 2013, 2021). Furthermore, according to the projected climatic conditions in a specific location and the analyzed crop-disease-climate system, different outcomes have been predicted for the future. Unchanged infection levels may be expected in the future, e.g. fire blight of apple trees in Switzerland (Hirschi et al., 2012) and late potato blight in Germany (Volk et al., 2010); in contrast, an increase of disease severity has been predicted for certain pathogens, due to more favorable future conditions, e.g. downy mildew and botrytis of grapevine (Salinari et al., 2006; Caffarra et al., 2012; Bregaglio et al., 2013; Gouache et al., 2011); climate change may instead decrease the infection risk, as predicted for black Sigatoka of banana plants (Jesus Júnior et al., 2008; Alves et al., 2011) and for Fusarium wilt of date palms (Shabani and Kumar, 2013). These studies focus on evaluating the association between different climatic factors and the considered pathogens to assess future disease pressure under different climate change scenarios, yet neglect the possible climate impacts on the plant host phenology and crop growth (but see Caffarra et al. (2012))

There is a need for a comprehensive and mechanistic understanding to disentangle the complex climate effects on multiple parasite and host traits in order to examine and predict the net effect of global warming on the considered pathosystem. The development and implementation of mathematical models has been identified as a key tool to overcome some of the described challenges related to climate change study and to guide the planning of adaptation strategies (Newbery et al., 2016; Juroszek and von Tiedemann, 2015). Multiple models (mechanistic, empirical and/or statistical) of infection dynamics (e.g. disease risk, yield loss and/or infected area) that take into account different environmental variables with diverse temporal resolutions and horizons have been implemented (Juroszek and von Tiedemann, 2015). Only few of the many available modeling approaches explicitly incorporate fundamental ecological processes that may determine the ability of a parasite species to respond to changing climate, such as rates of reproduction, survival and dispersal. This is the case of mechanistic/process-based models, which, compared to correlative models, are able to reconstruct the links between the functional traits of organisms and the relative environmental conditions (Leroux et al., 2013; Evans et al., 2015).

Although there has been a significant effort in developing mathematical models to examine the climate effects on the host plant dynamics (e.g. Vanalli et al. (2021); Chuine et al. (2016)) as well as on their pathogens (e.g. Bevacqua et al. (2023); Salinari et al. (2006)), modeling frameworks that link both impacts are quite uncommon (but see Caffarra et al. (2012)). Here, we aim at filling this gap, evaluating climate change impacts on crop production and disease-induced yield losses, mechanistically linking the climatic responses of the host plant and of its fungal pathogen. In this way, we can develop a generalizable modeling tool that can capture different climate change outcomes on the interacting components of the depicted disease triangle. To do so, we couple a phenological model we developed for blooming (subsequent chilling and forcing phases, see Vanalli et al. (2021)) and ripening (Growing Degree Day model, see Vanalli et al. (2021)) with a climate-driven model of Susceptible-Exposed-Infected type we develop for brown rot spread (Bevacqua et al., 2023). We consider key physiological and phenological processes of the plant (i.e. blooming, fruit ripening and growth) as well as the dynamics of the fungal pathogen (i.e. spore survival and infection) that are affected by temperature and precipitation. We use the peach-brown rot pathosystem at the national scale of France as a study case. The French metropolitan territory represents an excellent study case because i) it is the fourth biggest producer of peaches in Europe (*>*200,000 tons per year, see Bernadette and Bouchard-Aerts (2019)); ii) *M. fructicola* infection, the most common and widely distributed species, has been first detected in France, among the European countries, in 2001 (Lichou et al., 2002); iii) the French country is a good study system for evaluating climate change impacts, since it covers four climatic zones (Mediterranean, continental, oceanic and mountainous) in less than 600,000 km^2^.

Thus, we study and disentangle the ‘disease triangle’ of peach, brown rot and climate in France to evaluate historical baseline and future changes of production and yield loss due to brown rot infection and changing phenology for four groups of peach cultivars (i.e. early, mid-early, mid-late and late) under different scenarios of temperature and precipitation. Our predictions allow us to i) examine the long term trend of peach production at the country-level under climate change, ii) determine which regions are favorable not only for peach cultivation but also for producing a healthy yield, iii) identify critical areas, where brown rot spreading may be favored by future climatic conditions, and iv) maximize the national peach production by examining a possible adaptation strategy of shifting peach production sites to new suitable areas.

**Figure 1:**
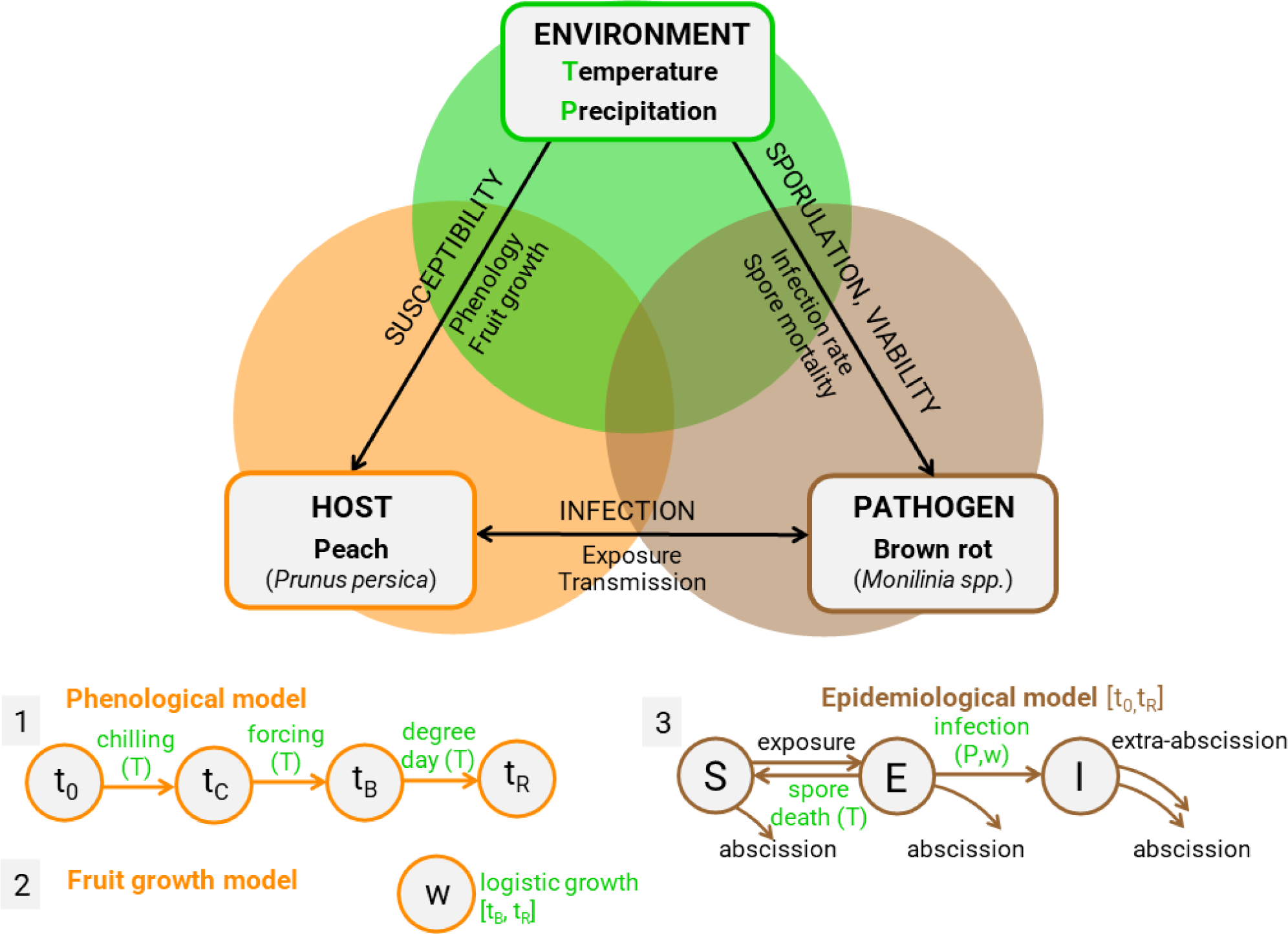
Schematic representation of the ‘disease triangle’ of brown rot infection (represented in brown) in peach orchards (in orange) under certain environmental conditions (temperature and precipitation, in green). The host phenological (1) and peach growth model (2) are specified, together with the type S-E-I epidemiological model (3). Temperature(T)- and precipitation(P)-dependent processes of each model are highlighted in green; more details are provided in the main text.

## 2 Methods

### 2.1 The plant host: peach phenology and fruit growth

The peach is the third most cultivated crop of the Rosaceae family (Obi et al., 2018) and has been extensively studied in both field and modeling works (Génard and Huguet, 1996; Ziosi et al., 2003; Allen et al., 2004; Lescourret et al., 2011), which established peach sensitivity to climate change (27; 63; 64). To simulate and predict future blooming and ripening times, we adopted a phenological modeling framework, which we have developed for four peach cultivars (see Vanalli et al. (2021); Chuine et al. (2016) for details). The temperature-driven phenological model consists of a subsequent chilling and forcing phase model for blooming time. The model assumes that time of endodormancy break (*t_C_*) occurs after accumulating enough chilling units. After, forcing units start to be accumulated up to a threshold that determines blooming time (*t_B_*). Blooming determines the beginning of the fruit growing season, which ends at harvest time when the fruit reach ripeness, according to a degree-day thermal model specified in (Vanalli et al., 2021). While different peach cultivars have similar thermal requirements for endodormancy and blooming, their parameters describing ripening and fruit growth greatly vary across different cultivars (Vanalli et al., 2021). Following Thornley and Johnson (1990), we assumed that a single fruit fresh mass (hereafter referred to as fruit weight) grows according to the Pearl-Reed logistic equation,

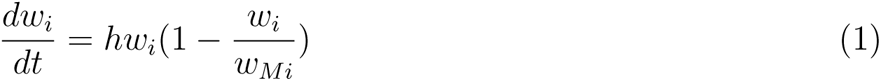

whose analytical solution in time is described by the following equation,

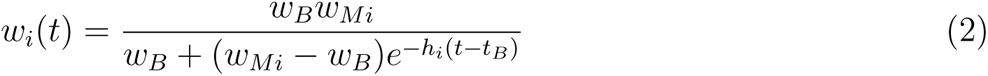

where *t* represents time (expressed in Day Of the Year, DOY), *i* represents the cultivar group (i.e. *i*=early, mid-early, mid-late, late cultivar), *t_B_*is the time at bloom, *w_B_*is the fruit weight at bloom in grams (g), *w_Mi_* is the cultivar-specific maximum fruit weight (g) and *h_i_* represents the conversion rate of resources into fruit mass (d^-1^) for the specific cultivar *i* (hereafter *h_i_* is referred to as fruit growth rate). We assumed that fruit weight at blooming time is the same across cultivars and that realized fruit weight at harvest (which is needed to compute the expected yield) corresponds to a fraction *C* of the potential fruit maximum weight for each cultivar (i.e. *w_Mi_* = *w_i_*(*t_Hi_*)*/C* = *w_Hi_/C*, see Appendix).

### 2.2 The fungal pathogen: brown rot

Brown rot is a fungal disease caused by *Monilinia* spp. which infects fruits and makes them rot, generating significant crop losses (Lichou et al., 2002; Larena et al., 2005). For this reason, it is considered as one of the most serious diseases of fruit orchards (Rungjindamai et al., 2014; Villarino et al., 2013). During the fruit growing season, after overwintering in twig cankers or mummified fruits or being transported by wind (Lacey, 1996; Holb, 2008; Byrde and Willetts, 1977; Aylor, 1999), *Monilinia* spp. conidial spores germinate and constitute the primary inoculum for infection. Primary inoculum spores are disseminated in the orchard and deposit on the fruit cuticles (Byrde and Willetts, 1977; van Leeuwen, 2000) where they can either die at a rate that is positively related to temperature (Xu et al., 2001a; Caubel et al., 2012; Bevacqua et al., 2023) or they can pass through the fruit cuticular surface via stomata, lenticels, wounds or cuticle cracks and successfully infect the fruit (Gibert et al., 2009; Xu et al., 2001b; Byrde and Willetts, 1977). The described infection process is facilitated by rain occurrence which increases the vulnerability of fruit cuticle to infection (Caubel et al., 2015; Bevacqua et al., 2023). The spores that successfully infect a fruit reproduce and generate a secondary inoculum that will cause new infections (Byrde and Willetts, 1977; Corbin, 1963), spreading throughout the orchard and causing consistent yield losses. We implemented a climate-driven epidemiological model to evaluate brown rot spreading during the fruit growing period, i.e. from blooming to ripening time, and the resulting disease-induced yield loss for each peach cultivar. This mechanistic compartmental SEI-like model has been developed with time-constant parameters by Bevacqua et al. (2018) using field experimental observations. Subsequently, Bevacqua et al. (2023) examined the climatic dependency of the multiple infection processes, selecting temperature *T* (*^◦^*C) and daily rain occurrence *P* (*P* = 0 if dry and *P* = 1 if wet) as significant drivers of spore mortality *η* and infection rate *σ*, respectively. Fruits are classified as susceptible (*S*), exposed to the pathogen (*E*) or infected and infectious (*I*). Their dynamics during the fruit growing period can be described by the following system of ordinary differential equations:

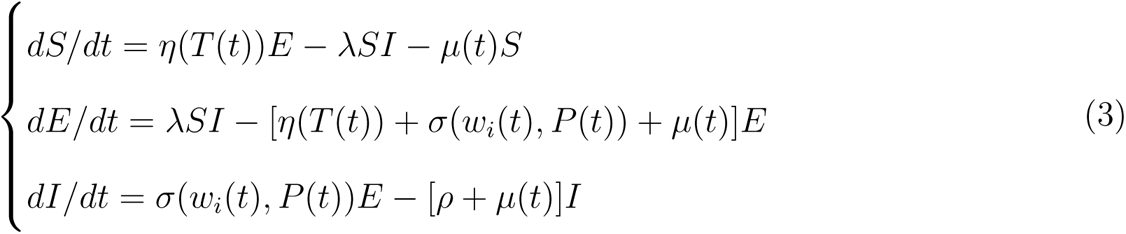

where *λ* is the exposure rate, *µ*(*t*) is the fruit age dependent peach abscission rate, and *ρ* is the extra abscission rate for infected peaches. Using the described model, for each cultivar and year, we computed the potential disease-free yield *Y*_0_, namely *S*(*t_H_*) obtained by the numerical integration of Eq. 2, with *E*(*t*) = *I*(*t*) = 0. The yield with disease *Y_d_* was computed as the sum of symptomless fruits at harvest time (*S*(*t_H_*) + *E*(*t_H_*), Eq. 3). Thus, the resulting percentage of brown-rot induced yield loss is calculated as 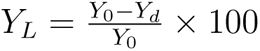. For the parameter values and the rationale justifying the above mentioned model formulation and assumptions, the reader can refer to the original model description in Bevacqua et al. (2018, 2023).

### 2.3 Environment: temperature and precipitation

Historical daily average and minimum air temperature (*^◦^*C) and precipitation occurrence time series have been made available by the project Drias (2013) developed by Météo-France (http://www.drias-climat.fr/. Accessed September 2023). The same project provided downscaled projections at a spatial resolution of 8 × 8 km^2^ (*∼* 0.11*^◦^* × 0.11*^◦^*) for the continental French territory (8,412 total cells) under an intermediate climatic scenario, Representative Concentration Pathway (RCP) 4.5, and a business as usual scenario, RCP 8.5 (IPCC, 2013) (https://www.drias-climat.fr/. Accessed September 2023). Climatic data are available in the historical period, from 1996 to 2015, while future projections under the two considered scenarios are from 2016 to 2100. The described models (Eq. 1,3) run with a daily time step, independently for each cell of the French metropolitan territory, from 1996 to 2100.

### 2.4 Adaptation scenario: spatial shifts of peach production sites

According to Vanalli et al. (2021), we assumed a cell to be suitable, in a given year, for peach cultivation if thermal requirements for endodormancy break, blooming and ripening are fulfilled. Water requirements for fruit growth and plant physiological processes are assumed to be fulfilled by irrigation. Because we showed a future spatial shift of the suitable areas for peach cultivation as a consequence of climate change (Vanalli et al., 2021), here, we considered two different scenarios: i) a no-adaptation scenario, i.e. peach cultivation areas for each cultivar are fixed to the estimated suitable historical regions (average of the decade 1996-2005) and ii) an adaptation scenario, i.e. peach cultivation areas are flexible to be shifted to new areas that were not suitable in the historical period (1996-2005) but that might become suitable in the future due to new climatic conditions. The comparison between the total yield obtained in these two scenarios allowed us to examine the efficacy of shifting peach production sites.

### 2.5 Peach yield at the national level

To provide a realistic estimate of the peach national yield, we examined the historical extension of peach orchards in the French continental territory, which consists of a total of 106 km^2^ (https://data.europa.eu/data/datasets. Accessed November 2023, (Radici et al., 2023)). We assumed that peach orchards are effectively cultivated only in those areas particularly suitable for peach cultivation, i.e. where any peach cultivar, form early to late maturing, can achieve ripening (Vanalli et al., 2021). In the considered historical period, the phenological model indicates that 390 (corresponding to 25,000 km^2^) of the 8,412 total spatial cells fulfills this condition. Thus, we assumed that: i) 0.27 km^2^ per “peach cultivation suitable” cell are devoted to peach orchards, ii) this extension is equally divided across the four cultivars, with 0.068 km^2^ per cell per cultivar, and iii) the remaining spatial cells, even if suitable for one or more cultivars, do not contribute to the peach national yield.

### 2.6 The interplay between the phenological and the epidemiological model

First, we examined the climate change impact on peach production considering only the climatic effect on peach phenology (i.e. in absence of brown rot) in the no-adaptation and in the adaptation scenarios. Second, we examined the climate change impact on peach production considering only the climatic effect on brown rot epidemiology while neglecting the climate-driven changes of peach phenology. In this case, peach phenology was kept fixed to the historical average in the decade 1996-2005. Last, we linked the direct climatic effects on plant phenology and fruit growth with the indirect climatic effects on brown rot dynamics, coupling the plant phenological model with the epidemiological model. In this way, we can disentangle the climate change impacts on the host and on the pathogen while examining their resulting synergism.

## 3 Results

Climate change impact on peach phenology We evaluated the impact of climate change on peach phenology and production in France under RCP 8.5 (Figure 2) and RCP 4.5 scenarios (Figure S1) in the absence of brown rot disease. High heterogeneity is expected across space and time and is dependent on the considered adaptation scenario.

**Figure 2:**
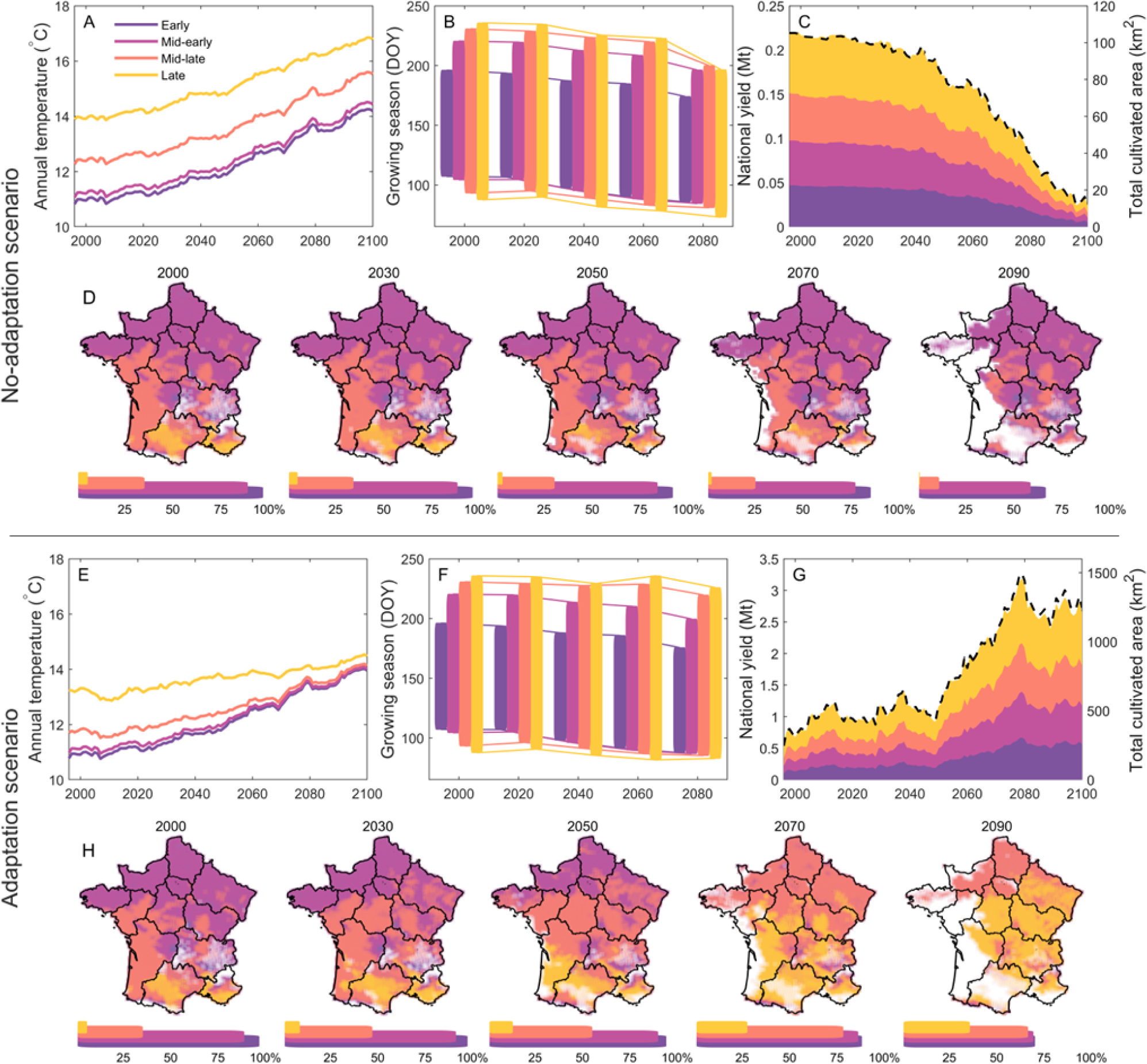
Climate change impact on peach phenology. Average annual temperature in Celsius degrees (A,E), average growing season expressed in Day Of the Year (DOY), in the decades 2000 (1996-2005), 2030 (2026-2035), 2050 (2046-2055), 2070 (2066-2075), 2090 (2086-2095) (B,F), national yield and its composition in Mega tons (left y-axis) and total cultivated area in km^2^ (black dashed line, right y-axis) (C,G), for early (dark purple), mid-early (bright purple), mid-late (orange) and late (yellow) cultivars under the RCP 8.5 climate scenario, in a no-adaptation (A-D) and in an adaptation (E-H) scenario of shift of peach production sites. Spatial distribution of suitable areas for peach cultivation in the decades 2000, 2020, 2040, 2060, 2080, with the relative percentage occupancy of the French territory (D_1_,_5_H); white spatial cells represent unsuitable areas for peach cultivation, i.e. null yield for more than one year over the decade. Lines in panels A,C,E,G are plotted with a moving average of 10 years to decrease inter-annual variability.

If the production sites for each cultivar remain the same as in the reference decade 1996-2005, we predict a drastic decline in peach production across the XXI century (Figure 2A-D). The estimated historical peach national yield from our model is consistent with the agricultural data reported by the European Commission of approximately 0.2 Mt (https://agriculture.ec.europa.eu/. Accessed November 2023). The average annual temperature is expected to increase in the future in a similar way across the French areas suitable for the four cultivars. Specifically, due to the high Degree-Day requirement for fruit ripening, late cultivars can be cultivated only in quite warm areas that are mainly situated in the south-western part of France and consist of less than 5% of the territory. In contrast, the less restrictive Degree-Day requirements for the mid-late, mid-early and early cultivars allow for their cultivation in colder and more extended regions (Figure2A,D). In the areas that will still be suitable for peach cultivation, increasing temperatures will accelerate the phenological cycle of all cultivars with an earlier blooming time and a faster ripening (Figure 2B). Conversely, in areas represented in white in Figure 2D, milder winter temperatures will fail the endodormancy chill unit requirement, impeding flowering and resulting in a null yield. These failure events will become significant at the end of the century and will mainly affect western France (2090, Figure 2C,D). This decline will cause a substantial drop in suitable areas for peach cultivation of less than 70% of the country, and there will be a nearly total disappearance of the suitable areas for late cultivars. Consequently, the potential French national peach yield will decline by almost 0.2 Mt from 2000 to 2100 (Figure 2C).

The alternative scenario, where peach cultivation sites are shifted to areas that were not suitable in the past, could overall have beneficial impacts, assuring a similar or larger peach yield in the future compared to historical production (Figure 2E-H). Thanks to the shift of production areas toward colder north-eastern regions, temperature conditions could be maintained close to the optimal range for plant phenological thermal requirements, i.e. cold temperatures in winter during endodormancy and warm temperatures in spring and summer for blooming and ripening (Figure 2E). Peach phenology will thus remain similar to the historical baseline (Figure 2F). The most striking difference of this adaptation scenario, compared to the no-adaptation scenario, is the expansion of the potential suitable areas for peach cultivation, and in particular for mid-late (+31%) and late cultivars (+30%) (Figure 2H), despite experiencing the same endodormacy break failures. As a consequence of the expansion of production sites, peach national yield is expected to increase with a higher contribution from later cultivars (Figure 2G).

Analyzing the same results for the RCP 4.5 scenario, we expect a more moderate temperature increase with fewer areas that will become unsuitable in the future (Figure S1). The adaptation scenario will still be beneficial but with a lower margin of yield increase (Figure S1).

Overall, climate change will strongly impact peach phenology and the consequent spatial distribution of the suitable areas for cultivation in France. This impact could result in a contraction of the peach production niche, with a decrease in the national peach yield. Otherwise, if the shift and/or expansion of production sites is possible, adaptation to climate change could be beneficial for the French economy and food security.

### 3.1 Climate change impact on brown rot epidemics

We evaluated the impact of climate change on brown rot epidemics and peach production in France under the RCP 8.5 (Figure 3) and RCP 4.5 scenarios (Figure S2), fixing plant phenological and fruit growth parameters (i.e. *t_B_*, *t*_0_, *t_R_*, and *h*) to the historical average in the decade 1996-2005 for each spatial cell. Temperature and precipitation changes during the peach growing season are expected to strongly affect brown rot outbreaks in the future.

**Figure 3:**
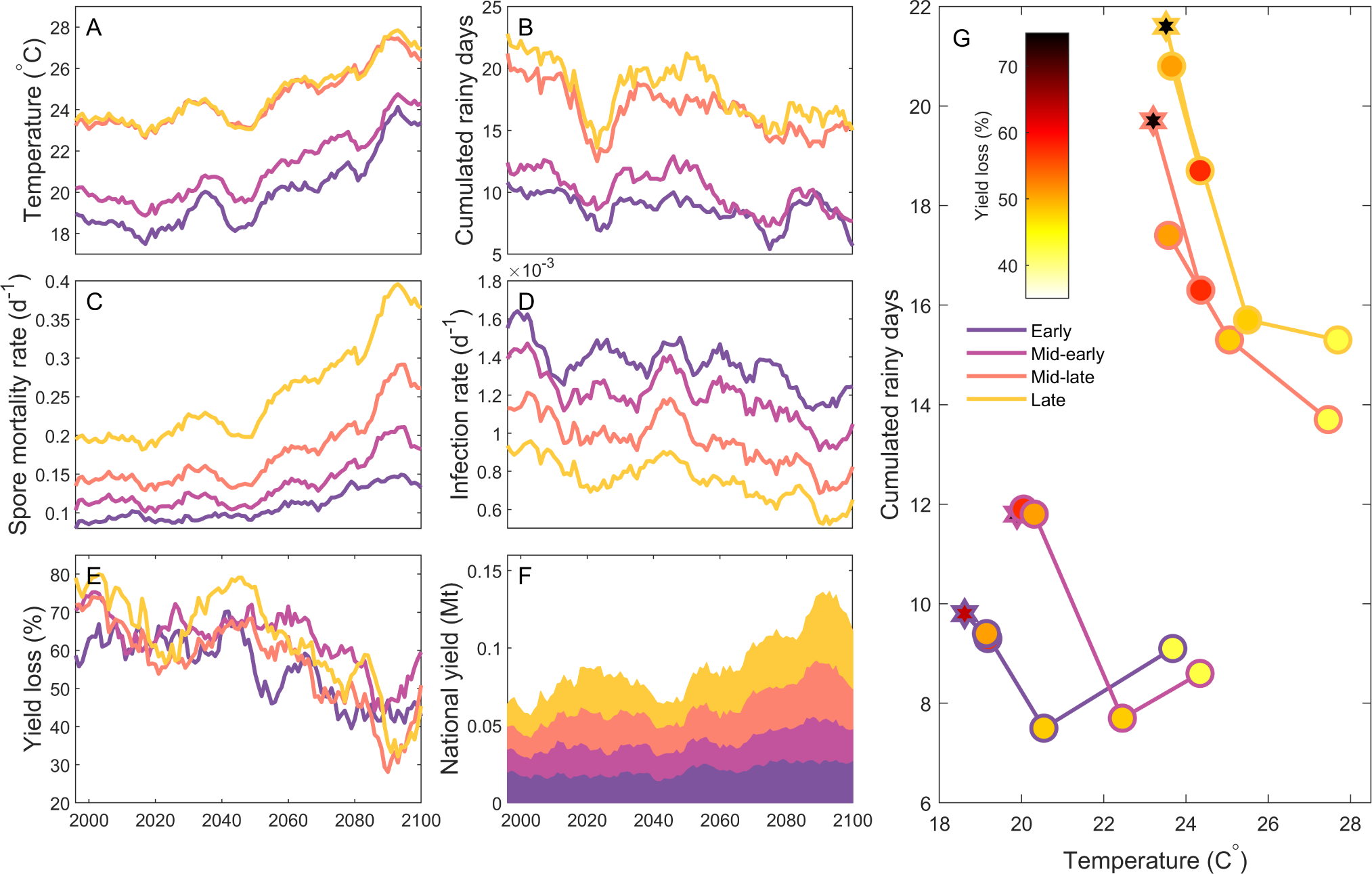
Climate change impact on brown rot disease. Average temperature during peach growing season (from blooming to harvest) in Celsius degrees (A), cumulated rainy days during peach growing season (B), fungal spore mortality rate (C), brown rot infection rate (D), yield loss due to brown rot compared to a disease-free yield (%) (E), and the resulting national yield (F) for early (dark purple), mid-early (bright purple), mid-late (orange) and late (yellow) cultivars, under the RCP 8.5 scenario. Lines in panels A-F are plotted with a moving average of 10 years to decrease inter-annual variability. The relation between temperature (x-axis), cumulated rainy days (y-axis) and yield loss due to brown rot (filling color) across the decades (2000 (star, 1996-2005), 2030 (2026-2035), 2050 (2046-2055), 2070 (2066-2075), 2090 (2086-2095)) is represented by connected dots for the four peach cultivars (outline color)(G).

Across the XXI century, climatic conditions during the peach growing season will be characterized by higher temperatures and fewer precipitation events (Figure 3A,B). This climatic trend will negatively affect disease spread due to a higher spore mortality rate and a lower infection rate (Figure 3C,D). For this reason, yield loss due to brown rot is expected to decrease, from nearly 70% to 40%, and national yield to increase, from 0.05 Mt to 0.13 Mt (Figure 3E,F). We note that the simulated national yield in presence of the disease is way lower compared to the statistics reported by the European Commission, because of the extensive use of fungicides which limits disease-induces losses (https://agriculture.ec.europa.eu/. Accessed November 2023). Examining the temporal trend of the peach yield losses due to brown rot for each cultivar in relation to temperature and cumulated rainy events, a lower yield loss across time (from dark red to white) in response to a warmer and drier climate is explicitly shown for each cultivar (Figure 3G). Analyzing the RCP 4.5 scenario disease projections, similar findings are highlighted for the future, but they are more moderate due to less extreme climatic conditions (Figure S2). Given the strict link between climatic conditions and brown rot spreading, climate change is expected to create less conducive conditions for disease spreading, resulting in less severe epidemics and higher healthy yield.

### 3.2 Synergistic impact of climate change on peach phenology and brown rot

We examined the synergistic impact of climate change on peach phenology and brown rot epidemics in France under RCP 8.5 (Figure 4) and RCP 4.5 scenarios (Figure S3), examining a no-adaptation and an adaptation scenario.

**Figure 4:**
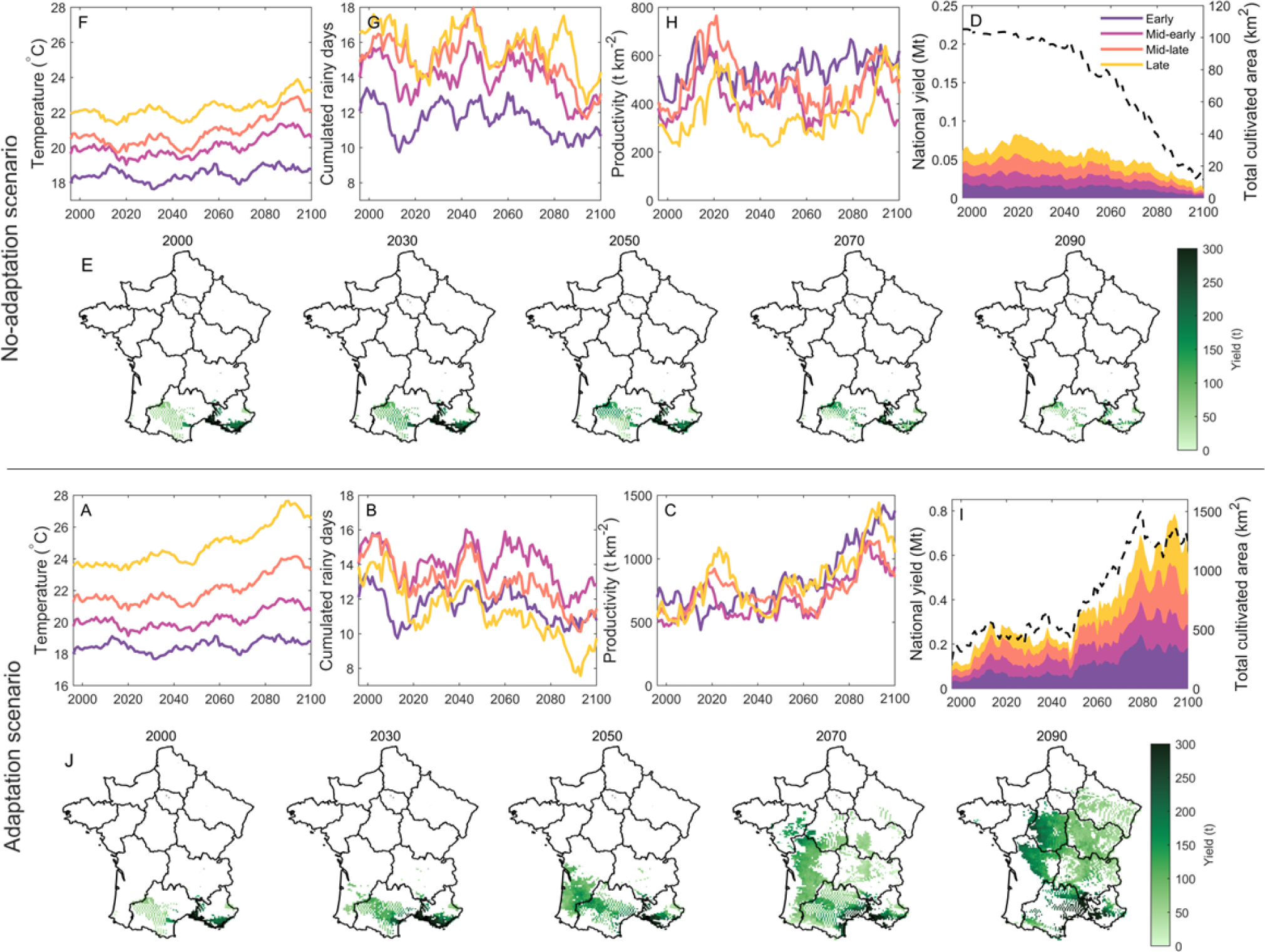
Synergistic climate change impact on peach phenology and brown rot disease. Average temperature during peach growing season (from blooming to harvest) in Celsius degrees (A,F), cumulated rainy days during peach growing season (B,G), yield productivity in tons per km^2^ (C,H), national yield and its composition in Mega tons (left y-axis) and total cultivated area in km^2^ (black dashed line, right y-axis) (D,I), for early (dark purple), mid-early (bright purple), mid-late (orange) and late (yellow) under the RCP 8.5 climate scenario, in a no-adaptation (A-E) and in an adaptation (F-J) scenario of shift of peach production sites. Lines in panels A-D and F-I are plotted with a moving average of 10 years to decrease inter-annual variability. Spatial distribution of total annual yield (from white to dark green) for peach cultivation in the decades 2000, 2030, 2050, 2070, 2090 (E,J); white spatial cells represent unsuitable areas for peach cultivation, i.e. null yield for more than one year over the decade.

In the no-adaptation scenario, peach production sites will experience higher temperatures and fewer rain events in the future (Figure 4A,B). Those conditions will disfavor brown rot spreading, with an increased productivity per km^2^, in particular for late cultivar (Figure 4C). Less severe brown rot epidemics will partially counterbalance the decline of suitable production sites at the end of the century, guaranteeing a comparable peach yield to the historical production (Figure 4D). The total peach yield varies significantly across space, due to how many cultivars can be cultivated and the severity of brown rot (Figure 4E). Historically (2000, decade 1996-2005), our simulations identify the south-eastern Mediterrenean area as the most productive. These regions coincide with the historical production sites for peach, namely Provence-Alpes-Ĉote d’Azur, Auvergne-Rĥone-Alpes and Occitanie. For the future, these areas will mostly turn unsuitable due to blooming failure caused by extreme warm conditions.

Under the adaptation scenario of shifting the peach production sites, temperatures will be less extreme and rain events will be more frequent than the previously described scenario (Figure 4F,G). Those climatic conditions are more conducive for brown rot spreading and will result in a lower peach productivity per km^2^ (Figure 4H). Despite this, there will still be an advantage in shifting the production areas, however, the potential benefit of such adaptation will be reduced due to more severe epidemics in these new regions (Figure 4I). Independently from the considered adaptation scenarios, the most productive areas for peach cultivation at the end of the XXI century will not be located anymore in the south-western area but in the central part of the country.

Examining the same projections for the more moderate RCP 4.5 scenario, we found similar results comparing the no-adaptation with the adaptation scenario (Figure S3). Independently from adaptation, it is worth noticing that the the Mediterranean region will still produce the highest peach yield at the end of the century, together with a new region situated in the north-western part of the territory (Figure S3).

In conclusion, moving peach production sites to new areas could potentially increase peach production, however, climatic conditions in those new regions could facilitate disease spreading and result in a lower net yield.

### 3.3 Effectiveness of adaptation: Spatial shift of peach production sites

We estimated the effects in time on yield loss due to brown rot disease and on the total national yield of shifting the peach production sites as a possible adaptation strategy to climate change (Figure 5).

**Figure 5:**
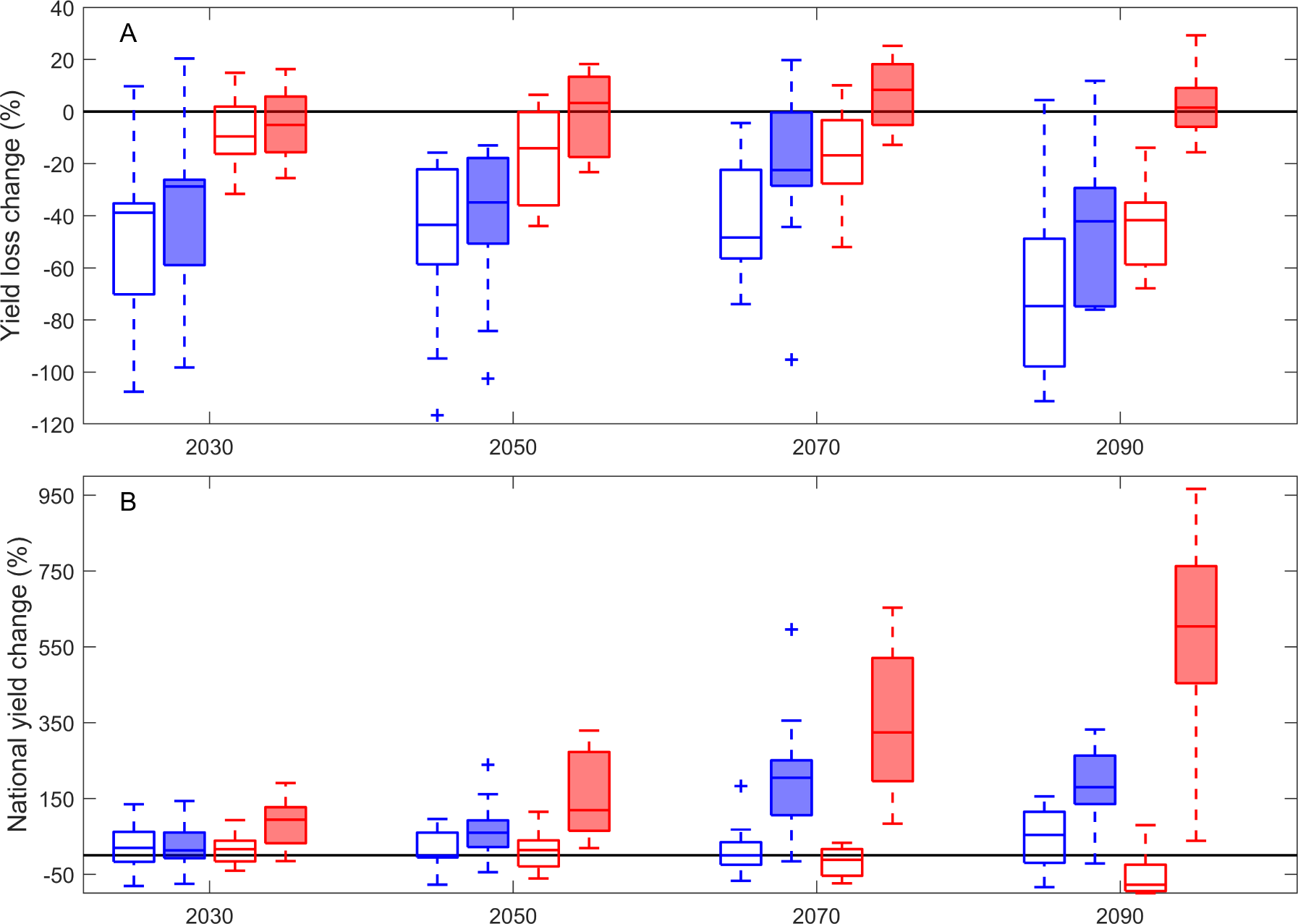
Effectiveness of adaptation for peach production. Boxplots of percentage variations of yield loss due to brown rot (A) and national yield (B) compared to the historical average in the decade 1996-2005 in the years 2020 (2016-2025), 2040 (2036-2045), 2060 (2056-2065), and 2080 (2076-2085)), for the RCP 8.5 (in red) and RCP 4.5 (in blue) climatic scenarios, with (filled boxplots) and without adaptation (empty boxplots). The black line in each panel highlights the null change, 0%, compared to historical levels.

Adaptation will result in slightly higher disease-induced yield losses compared to the no-adaptation scenario due to the more favourable conditions for disease spreading (Figure 5A,3). Under the RCP 8.5 scenario, disease losses will be comparable to those experienced historically, while we expect a significant decrease of yield losses up to -70% under the RCP 4.5 scenario. A consistent higher national yield for both the RCP 4.5 and RCP 8.5 scenarios is expected in the long-term future and in particular in the adaptation scenario, due to the spatial expansion of peach production sites (Figure 5B).

Even though worse disease outbreaks are expected when we considered adaptation, the overall yield would benefit from shifting and expanding peach production areas in both an indeterminate (RCP 4.5) and a business as usual (RCP 8.5) scenario.

## 4 Discussion

In this study, we examined the climate change impacts on peach phenology and brown rot disease across the XXI century in continental France under a moderate and an extreme climate change scenario. Specifically, we built on previous knowledge on the climatic responses of plant phenology (Vanalli et al., 2021) and of brown rot spread (Bevacqua et al., 2023), bridging these climate-dependent processes that were previously studied independently and we have now evaluated the entire ‘disease triangle’ of brown rot infection in peach orchards. We found that the suitable areas for peach cultivation will be drastically shifted by the end of the century for the RCP 8.5 scenario and less so for the RCP 4.5 scenario due to a combination of chilling and forcing requirements, with the first being hardly met in the future and the latter more easily fulfilled. In the case where the peach production sites are not shifted, an increase in peach productivity per unit of area will be experienced because of less severe brown rot outbreaks in a drier and warmer climate. However, due to too mild winters that are responsible for failing the peach endodormancy requirements, the suitable areas for peach production will drastically decline. Alternatively, we evaluated the spatial shift of production sites to new suitable areas as a form of adaptation. This strategy is projected to expand peach production to new areas and result in a net higher national yield. However, our simulations indicate that in those new regions, climate conditions will be conducive for brown rot spreading and proper disease control measures might be needed.

Understanding and foreseeing the impacts of climate change on crop production allows to inform and optimize adaptation measures to limit the negative expected impacts. The adaptation strategy that we considered in this work is the relocation of peach production sites within the French borders. Although this strategy is of complex implementation, depending on the climate change scenario that will be realized in the future, a spatial planning of planting sites might be necessary to safeguard future yield (Läderach et al., 2017; Ovalle-Rivera et al., 2015). In this case, a strategic spatial planning that incorporates land use change when evaluating the potential shift of suitable areas for peach cultivation is required. Importantly, orchard cultivation is not only a mere economic and food production sector but guarantees important ecosystem services, including climate and water regulation, soil nitrogen availability, and pollination (Demestihas et al., 2017). Furthermore, Mediterranean orchard-dominated landscapes are part of the cultural heritage of this territory, which is essential for both the local communities and tourists. Counter to this, the introduction of such landscapes in new regions for climate adaptation could represent an unwanted novelty (Montanaro et al., 2017) and might face territory ownership issues and pedological restrictions. These practical, agricultural, ecological and cultural aspects should not be overlooked.

Given the contrasting climatic requirements and the different susceptibility to diseases of different cultivars of the same crop, cultivar selection represents an additional advantage for farmers to adapt to a changing climate and guarantee a future yield. Even though the cultivar adaptation has been identified as one of the most effective agricultural strategies (Challinor et al., 2014), the benefits of such a measure across different regions, temporal horizons and projected climate change scenarios still need further investigation Zabel et al. (2021). In this work, we showed the different responses between early, mid-early, mid-late, and late cultivars, and we identified which cultivar, for each spatial cell of France, is expected to i) produce a potential higher yield and ii) have the least loss from brown rot. Our findings indicate that the choice of selecting an early cultivar, which has a shorter growing season and is less susceptible to brown rot infection, could be preferred to a later cultivar in hot-spot sites for brown rot disease. In contrast, in areas where climatic conditions disfavor brown rot spreading, the use of later cultivars, which have a higher fruit size at harvest, might be more profitable. The cultivar shift as a form of adaptation to climate change has been proposed and/or implemented for other important crop systems, such as rice, maize and grapes (Lv et al., 2020; Ausseil et al., 2021; Yoshida et al., 2015). However, this adaptation strategy has been based on the phenological requirements of the plant host and has rarely included the effect of pathogens and pests on the effective yield. Even more, there is the potential for the genetic improvement and development of new cultivars with low chilling requirements and/or that are more resistant to pathogens and extreme climatic events (Zhang et al., 2018).

Adaptation measures are not the only option to safeguard future crop yield. Developing more effective disease control strategies also represents a key resource to produce a healthy yield. However, plant disease control is heavily dependent on pesticides and fungicides to fight the variety of pathogens and pests that threaten crop production (Lucas et al., 2015). Despite their effectiveness in disease management, those strategies have important drawbacks. The emergence of resistance to many of the most effective fungicides has been compromising their efficacy (Lucas et al., 2015). Furthermore, there are some concerns related to chronic mammalian toxicity and environmental pollution from the chemical compounds of the fungicides (Waard et al., 1993). Therefore, the application of fungicides does not seem to be the long-term optimal solution to safeguard crop production and profitability. In addition, there is a growing demand of organic goods on the market, which prohibits the use of synthetic fertilizers and pesticides, while enhancing biodiversity and soil quality (Reganold and Wachter, 2016). This reinforces the need for the development and implementation of mathematical models that aim to understand climate change impacts on crop production and inform adaptation and control strategies to move agriculture towards a more sustainable and efficient system that can rely less on the use of chemicals for disease control.

In this work, we evaluated a simplification of the diverse range of possible adaptation and control measures that could be implemented in the future to safeguard peach production in the French territory. Specifically, we assumed an equal extension between the four considered groups of cultivars for a suitable cell for peach cultivation. This assumption is a simplification of the potential spatial heterogeneity that could arise in those areas due to specific cultivar choices by farmers, which could also change in the future. Moreover, even though in the considered adaption scenario we predicted an expansion of peach cultivation areas, this might subtract suitable agricultural land to other types of crops. Additional socioeconomic factors will indeed affect whether and how such expansion would be implementable. Additionally, the inclusion of other climate change scenarios could increase the strength of our findings. Here, we examined an intermediate (RCP 4.5) and a more extreme (RCP 8.5) emission scenario, with the latter being lately classified as too extreme and unlikely (Hausfather and Peters, 2020).

## 5 Conclusions

Our study provides new insights into the understanding of climate change impacts on crops, examining the two fold effects of climate on the plant host as well as on the fungal pathogen. The mechanistic mathematical framework implemented here is generalizable across cultivars or can be applied to other important fruit crop systems for the French economy. Whether and how will climate change impact peach production in France? Our findings indicate that global warming is expected to severely impair fruit production and modify brown rot infection dynamics, however, the implementation of adaptation strategies, such as the spatial shift of peach production sites could limit the expected losses. Predictive modeling frameworks like ours, which tackle the impacts of climate change on agricultural systems, provide key tools to face the great challenges of the future, such as food security, global warming and infectious diseases.

## Acknowledgments

We would like to acknowledge Dr. Isabella Cattadori and Dr. Davide Martinetti for their valuable support. Beth Tuschhoff is also thanked for her help in proofreading this paper.

