## Appendix for "Phenological and epidemiological impacts of climate change on peach production"

### Appendix A: Supplementary material

We assumed the potential fruit maximum weight for each cultivar, $w_{Mi}$, proportional to the fruit weight at harvest, $w_i(t_{Hi})$, according to a constant $C$. From Bevacqua et al. Bevacqua et al. (2018), we computed $C = 77\%$, and we retrieved realized cultivar-specific fruit weight at harvest from published literature studies (see Bussi et al. (1995); Zarzar et al. (2020); Selli and Bassi (1990) for early-cultivar, Bussi et al. (2002); Remorini et al. (2008) for mid-early cultivar, Öhlinger et al. (2008) for mid-late cultivar and Li et al. (1989); Fallahi et al. (2009) for late cultivar). In this way, for each year, given the climatic dynamics in each spatial cell, it is possible to calculate the blooming time $t_B$, the harvest time of each cultivar $t_{Hi}$ and the cultivar-specific maximum fruit weight $w_{Mi}$. Then, it is possible to estimate the fruit growth rate $h_i$, inverting the logistic equation of fruit growth (Eq 1,2). Note that the fruit growth rate inversely depends on the duration of the growing season $t_{Hi} - t_B$, which is affected by temperature conditions all year round.

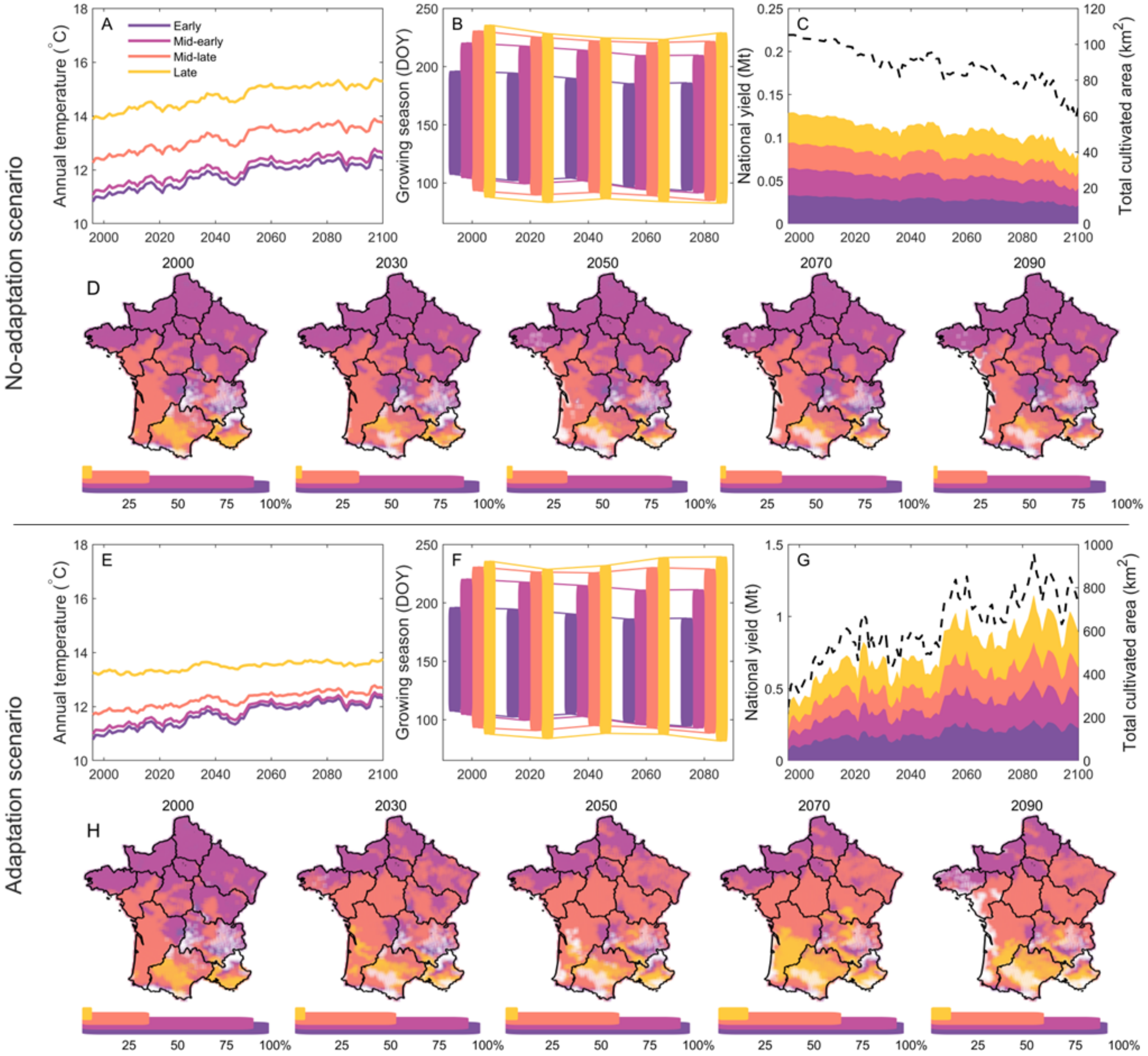


Figure S1: **Climate change impact on peach phenology.** Average annual temperature in Celsius degrees (A,E), average growing season expressed in Day Of the Year (DOY), in the decades 2000 (1996-2005), 2030 (2026-2035), 2050 (2046-2055), 2070 (2066-2075), 2090 (2086-2095) (B,F), national yield and its composition in Mega tons (left y-axis) and total cultivated area in km$^2$ (black dashed line, right y-axis) (C,G), for early (dark purple), mid-early (bright purple), mid-late (orange) and late (yellow) cultivars under the RCP 4.5 climate scenario, in a no-adaptation (A-D) and in an adaptation (E-H) scenario of shift of peach production sites. Spatial distribution of suitable areas for peach cultivation in the decades 2000, 2020, 2040, 2060, 2080, with the relative percentage occupancy of the French territory (D,H); white spatial cells represent unsuitable areas for peach cultivation, i.e. null yield for more than one year over the decade. Lines in panels A,C,E,G are plotted with a moving average of 10 years to decrease inter-annual variability.

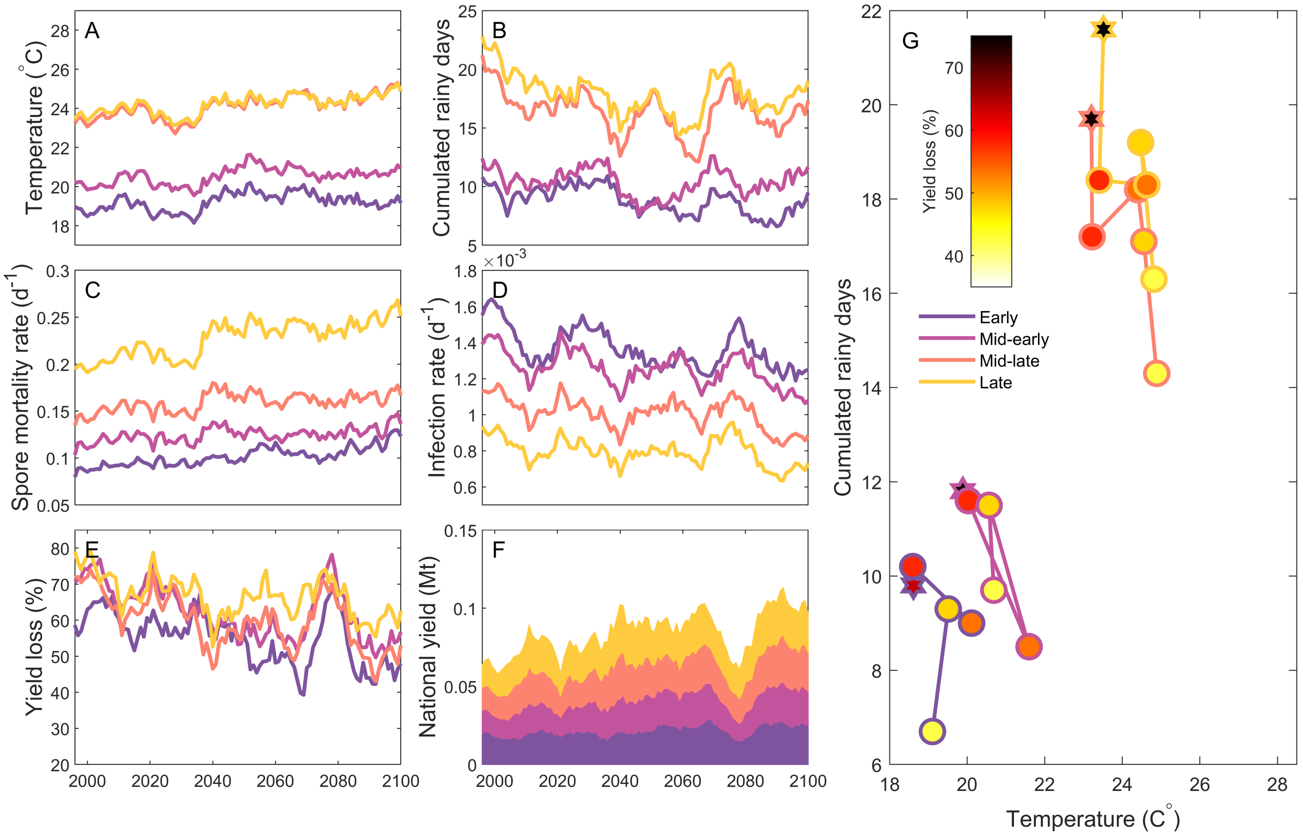


Figure S2: **Climate change impact on brown rot disease.** Average temperature during peach growing season (from blooming to harvest) in Celsius degrees (A), cumulated rainy days during peach growing season (B), fungal spore mortality rate (C), brown rot infection rate (D), yield loss due to brown rot compared to a disease-free yield (%) (E), and the resulting national yield (F) for early (dark purple), mid-early (bright purple), mid-late (orange) and late (yellow) cultivars, under the RCP 4.5 scenario. Lines in panels A-F are plotted with a moving average of 10 years to decrease inter-annual variability. The relation between temperature (x-axis), cumulated rainy days (y-axis) and yield loss due to brown rot (filling color) across the decades (2000 (star, 1996-2005), 2030 (2026-2035), 2050 (2046-2055), 2070 (2066-2075), 2090 (2086-2095)) is represented by connected dots for the four peach cultivars (outline color) (G).

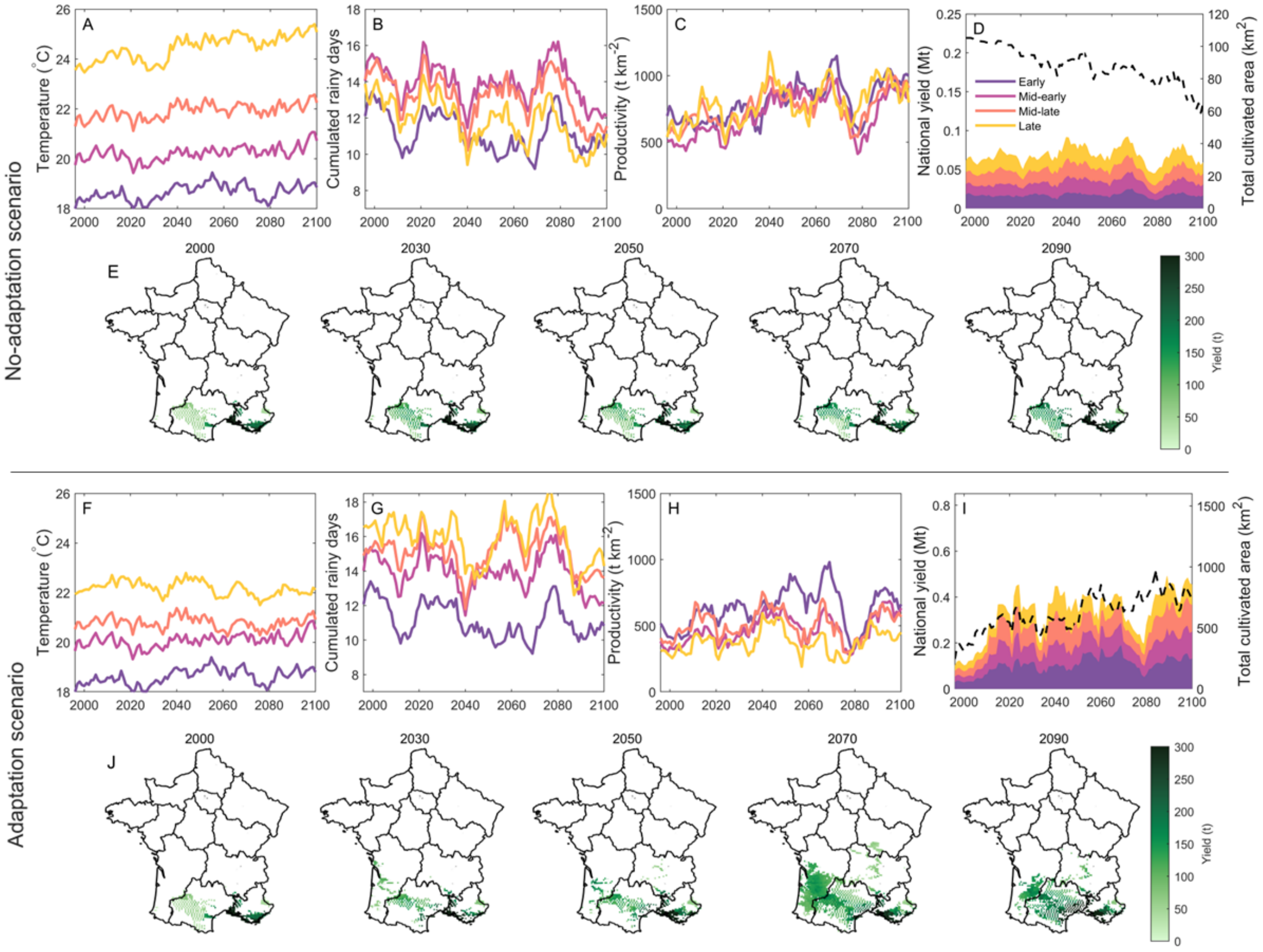


Figure S3: **Synergistic climate change impact on peach phenology and brown rot disease.** Average temperature during peach growing season (from blooming to harvest) in Celsius degrees (A,F), cumulated rainy days during peach growing season (B,G), yield productivity in tons per km$^2$ (C,H), national yield and its composition in Mega tons (left y-axis) and total cultivated area in km$^2$ (black dashed line, right y-axis) (D,I), for early (dark purple), mid-early (bright purple), mid-late (orange) and late (yellow) under the RCP 4.5 climate scenario, in a no-adaptation (A-E) and in an adaptation (F-J) scenario of shift of peach production sites. Lines in panels A-D and F-I are plotted with a moving average of 10 years to decrease inter-annual variability. Spatial distribution of total annual yield (from white to dark green) for peach cultivation in the decades 2000, 2030, 2050, 2070, 2090 (E,J); white spatial cells represent unsuitable areas for peach cultivation, i.e. null yield for more than one year over the decade.

Rania Zarzar, Abdelali Blenzar, Adil Asfers, Abelhadi Ait Houssa, and Ossama Kodad. Year effect on blooming and harvesting time and fruit weight in two peach varieties grown under sais valley climatic conditions. In *E3S Web of Conferences*, volume 183, page 03003. EDP Sciences, 2020.
